# Compartmentalized control of Cdk1 drives mitotic spindle assembly

**DOI:** 10.1101/2021.04.20.440650

**Authors:** Angela Flavia Serpico, Francesco Febbraro, Caterina Pisauro, Domenico Grieco

## Abstract

During cell division, dramatic microtubular rearrangements driven by cyclin B-cdk1 (Cdk1) kinase activity mark mitosis onset leading to interphase cytoskeleton dissolution and mitotic spindle assembly. Once activated by Cdc25, that reverses inhibitory phosphorylation operated by Wee1/Myt1, Cdk1 clears the cytoplasm from microtubules by inhibiting microtubule associated proteins (MAPs) with microtubule growth-promoting properties. Nevertheless, some of these MAPs are required for spindle assembly, creating quite a conundrum. We show here that a Cdk1 fraction bound to spindle structures escaped Cdc25 action and remained inhibited by phosphorylation (i-Cdk1) in mitotic human cells. Loss or restoration of i-Cdk1 inhibited or promoted spindle assembly, respectively. Furthermore, polymerizing spindle microtubules fostered i-Cdk1 by aggregating with Wee1 and excluding Cdc25. Our data reveal that spindle assembly relies on compartmentalized control of Cdk1 activity.

---

To prevent chromosome damage during spinlde assembly, the microtubular interphase cytoskeleton must be dismantled at the onset of mitosis and, as many mitotic features, this is an event downstream Cdk1 activation. Indeed, Cdk1 activity clears cytoplasmic microtubules by directly or indirectly affecting several microtubule-stabilizing and -destabilizing MAPs, inhibiting, for instance, the microtubule growth-promoting properties of ch-Tog and Map4 (*1, 2, 3, 4, 5*). Nevertheless, a number of these MAPs are required for spindle microtubule growth (*2, 6, 7*). Thus, how spindle assembles despite the potent microtubule-destabilizing action of Cdk1 is unknown. During interphase, Cdk1, accumulates in an inactive state, phosphorylated at inhibitory sites by Wee1/Myt1 kinases (*8*). At mitosis onset, initial reversal of these phosphorylation by Cdc25 phosphatase is rapidly followed by full Cdk1 activation via positive feedback loops in which Cdk1 inhibits Wee1/Myt1 and stimulates Cdc25 (*8*). Thus, inhibitory phopshorylation of Cdk1 has a crucial role in ensuring completion of interphase tasks before mitosis onset, but once Cdk1 activation begins this control appears to be rapidly suppressed by autoactivatory feedback loops. Cdk1 is believed to be only inactivated upon spindle assembly by cyclin B degradation (*8*). Nevertheless, early evidence indicated that cells expressing a Cdk1 version non-phosphorylatable at the inhibitory sites threonine 14 and tyrosine 15 (Cdk1AF), rapidly entered mitosis but failed to assemble spindles (*9*). These mitotic defects were, however, ascribed to premature mitosis onset in presence of incompletely replicated DNA and not as a requirement for inhibitory phosphorylation of Cdk1 in spindle assembly (*9*). In addition, evidence that i-Cdk1 is present in meiotic and mitotic extracts from Xenopus eggs and mouse eggs was later reported (*10, 11*). Here we examined if and how inhibitory phosphorylation of Cdk1 had a role for spindle assembly.

To this end, we begun analysing spindle assembly upon downregulation of Wee1 expression by small interfering RNAs (siRNAs) in hTERT-RPE1 cells. To prevent premature mitosis onset upon Wee1 downregulation, 6 hours post Wee1-siRNA treatment cells were incubated with the selective and reversible Cdk1 inhibitor RO-3306 at 9 μM (high-RO) for further 16 hours, to allow completion of DNA replication and arrest cell cycle at G2 (*12*). As control, part of Wee1-siRNA-treated cells were previously complemented with siRNA-resistant Wee1 expression vector (fig. 1A; Fig. S1) (*13*). Upon release from G2-arrest into fresh medium, containing the proteasome inhibitor MG-132 to block mitosis exit and the protein synthesis inhibitor cycloheximide to prevent abnormal protein accumulation (M/C medium), the majority of control cells built normal bipolar spindles within 60 min incubation, that remained stable for further 20 min incubation (fig. 1A). Conversely, spindle assembly was markedly impaired in Wee1-downregulated cells, with defects that ranged from monopolarity to poorly structured bipolar spindles with gross chromosome alignment abnormalities (fig. 1A). To establish whether impaired spindle assembly was due to Wee1 loss *per se* or because it affected Cdk1 activity, we asked if mild Cdk1 inhibition could restore spindle assembly in Wee1-downregulated cells. Indeed, addition of RO-3306 at 0.5 μM (low-RO), from 60 to 80 min post G2 release, substantially restored spindle assembly (fig. 1A). Defects in spindle assembly were also induced by cdk1AF overexpression under similar conditions of G2-arrest and release, paralleling what described by Krek and Nigg in 1991, as well as in Wee1-siRNA-treated HeLa cells or by chemical inhibition of Wee1, in all cases defects were substantially reversed by low-RO (figs. S2, S3 and S4A) (*9, 14*). In another set of experiments, non-targeting-siRNA-, as control, and Wee1-siRNA-treated hTERT-RPE1 cells were arrested at prometaphase by the reversible microtubule inhibitor nocodazole added shortly after siRNA treatments (as described in Materials and Method). Cells were, then, released in M/C medium for 80 min and a portion of Wee1-siRNA-treated cells also received low-RO at 60 min incubation. Within this period, the majority of control cells assembled spindles, while assembly was strongly impaired in Wee1-siRNA-treated cells but restored by low-RO (fig. 1B). When cells were released just into fresh medium, mitosis exit was delayed in Wee1-siRNA-treated cells, as shown by delayed cyclin B1 degradation, presumably by the action of the spindle assembly checkpoint (SAC; fig. S5) (*15*). Together, these data suggest that inhibitory phosphorylation-dependent control of Cdk1 activity is required for spindle assembly.

**Fig. 1.**
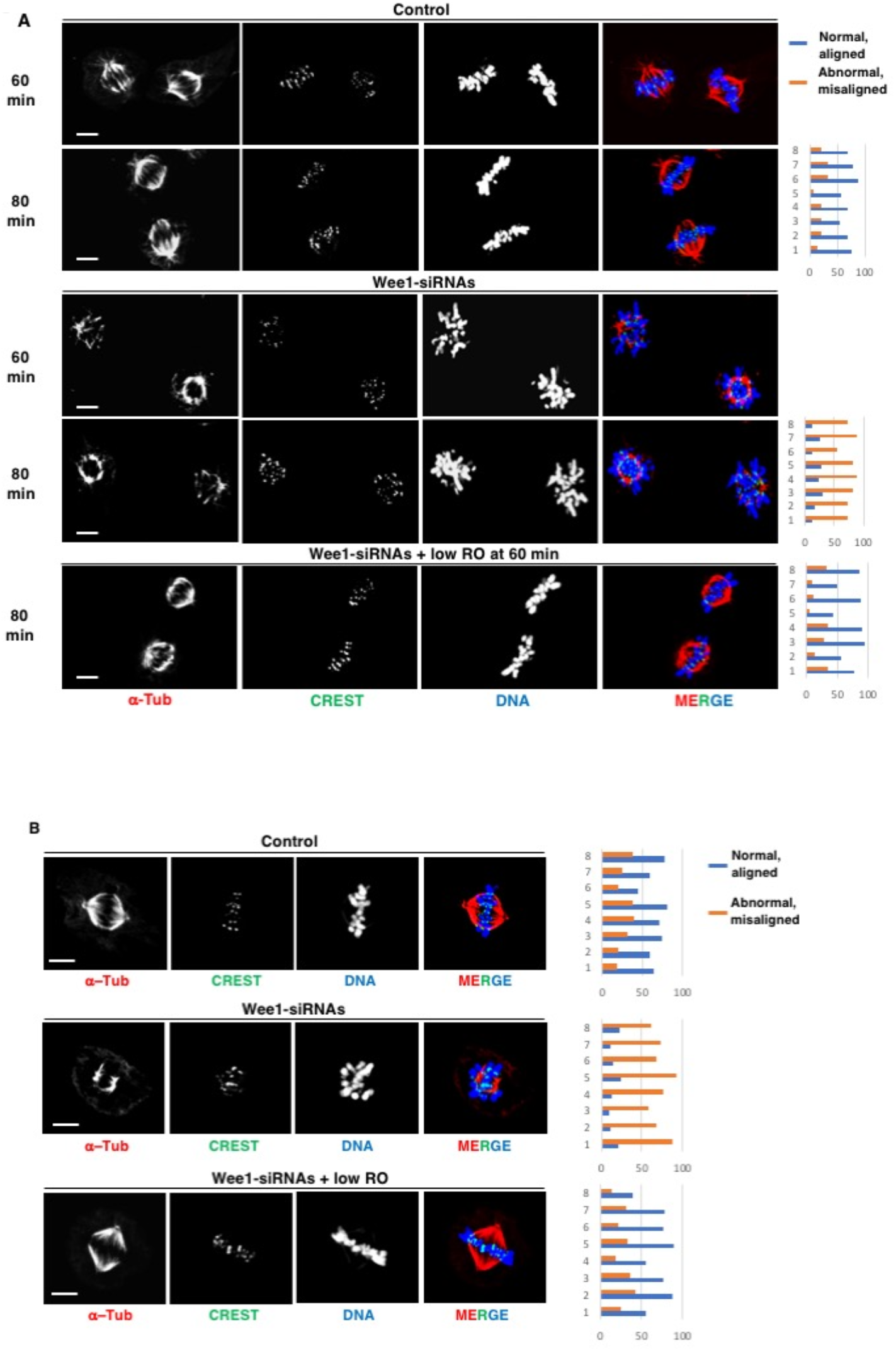
Wee1 downregulation affects spindle assembly. (**A**) hTERT-RPE1 cells were arrested at G2 by high-RO treatment, released and fixed at 60 and 80 min of incubation in M/C medium (see Methods section). Control: Wee1-siRNA-treated cells previously transfected with siRNA-resistant Wee1 expression vector; Wee1-siRNAs: Wee1-siRNA-treated cells previously transfected with empty vector; Wee1-siRNAs + low-RO: Wee1-siRNAs to which low-RO was added by 60 min from G2-arrest release. (**B**) hTERT-RPE1 cells were arrested at prometaphase with nocodazole, released and fixed after 80 min of incubation in M/C medium (see Materials and Methods). Control: cells treated with non-targeting siRNAs; Wee1-siRNAs: just Wee1-siRNA-treated cells; Wee1-siRNAs + low-RO: Wee1-siRNA-treated cells to which low-RO was added by 60 min from prometaphase-arrest release. Cells were fixed and stained for CREST (centromere marker, green), α-tubulin (a-Tub, red) and DNA (blue). Addition of vehicle (DMSO) had no effect on spindle assembly in Wee1-siRNA-treated cells. Graphs: quantitation of cells containing normal aligned (blue bar) or abnormal misaligned (orange bar) spindles at 80 min incubation; shown are data from two independent experiments (1-4, experiment 1; 5-8, experiment 2), around 100 cells were scored in 4 independent microscopy slide fields. Scale bar: 5 μm.

To directly test if i-Cdk1 was present in mitotic cells and where it was localised, we probed metaphase cells with an antibody recognizing cdk1 phosphorylated at tyrosine 15 (p-Y15-cdk1; fig. 2A). The p-Y15-cdk1 signal appeared decorating the spindle area paralleling total cdk1 localization (fig. 2A). To biochemically dissect localization and function of mitotic i-Cdk1, we adapted a method described by Nigg and coworkers to isolate mitotic spindles and tightly associated proteins by separating microtubular, insoluble, pellet fraction (P) from soluble fraction (S) of mitotic cells (see Materials and Methods) (*16*). Nocodazole-treated, prometaphase-arrested hTERT-RPE1 cells (Pro) were collected and taken at time 0 or taken after release in M/C medium for 60 min to have metaphase-arrested cells (Meta; fig. 2B). Cells were fractionated and proteins from S and P fractions analyzed (fig. 2B; P fraction was enriched 4 folds relatively to S fraction). In Pro cell P fraction, centrosomes were present as indicated by significant amounts of the central protein γ-tubulin and Nuclear and Mitotic Apparatus protein 1 (NuMA1), a centrosome-associated protein (fig. 2B) (*3*). Conversely, α-tubulin was substantially present in S fraction and much less in P fraction, as expected because of the microtubule polymerization block, as well as the MAPs Map4 and ch-Tog (fig. 2B). In Meta cells, instead, α-tubulin and significant amounts of Map4 and ch-Tog, the latter also involved in activating centrosome-independent, intra-spindle, microtubule nucleation, were present in P fraction, reflecting spindle assembly (fig. 2B) (*17, 18*). Distribution of NuMA1 and γ-tubulin in S and P fractions did not substantially change between Pro and Meta cells (fig. 2B). P fraction also contained small amounts of Wee1 that increased in Meta cells, while minimal amounts of Cdc25C were equally present in P fraction of both Pro and Meta cells (fig. 2B; similar protein distribution was found in HeLa cells under similar experimental conditions; see Fig. S6). Small amounts of cyclin B1 and cdk1 were present in P fraction of Pro cells and more of Meta cells (fig. 2B). However, i-Cdk1 was found quite exclusively in P fractions, as shown by p-Y15-cdk1 signal and by retarded cdk1 mobility on SDS/PAGE, and more from Meta than from Pro cells (Fig. 2B) (*19*). Quantitation of the upshifted cdk1 forms from Meta cell P fraction indicated that i-Cdk1 represents less than 10% of total Cdk1. The selective presence of i-Cdk1 in P fraction was confirmed also by probing immunoprecipitates (Ips) of comparable amounts of cyclin B1 from S and P fraction of Meta cells (Fig. 2C). In Meta cell P fraction, Map4 and ch-Tog appeared to have increased mobility on SDS/PAGE compared to S fraction, suggesting that these proteins were relatively dephosphorylated when microtubule-bound. Indeed, when Meta cells were further treated with nocodazole, from 60 to 80 min, to revert to a prometaphase condition, distribution of proteins between S and P fractions returned similar to Pro cells; Map4 and ch-Tog regained slower migration on SDS/PAGE, presumably reflecting their re-phosphorylation when returning into the cytoplasm, while Wee1 and i-Cdk1 were reduced in the P fraction, indicating their dependence also on microtubule polymerization (fig. 2D).

**Fig. 2.**
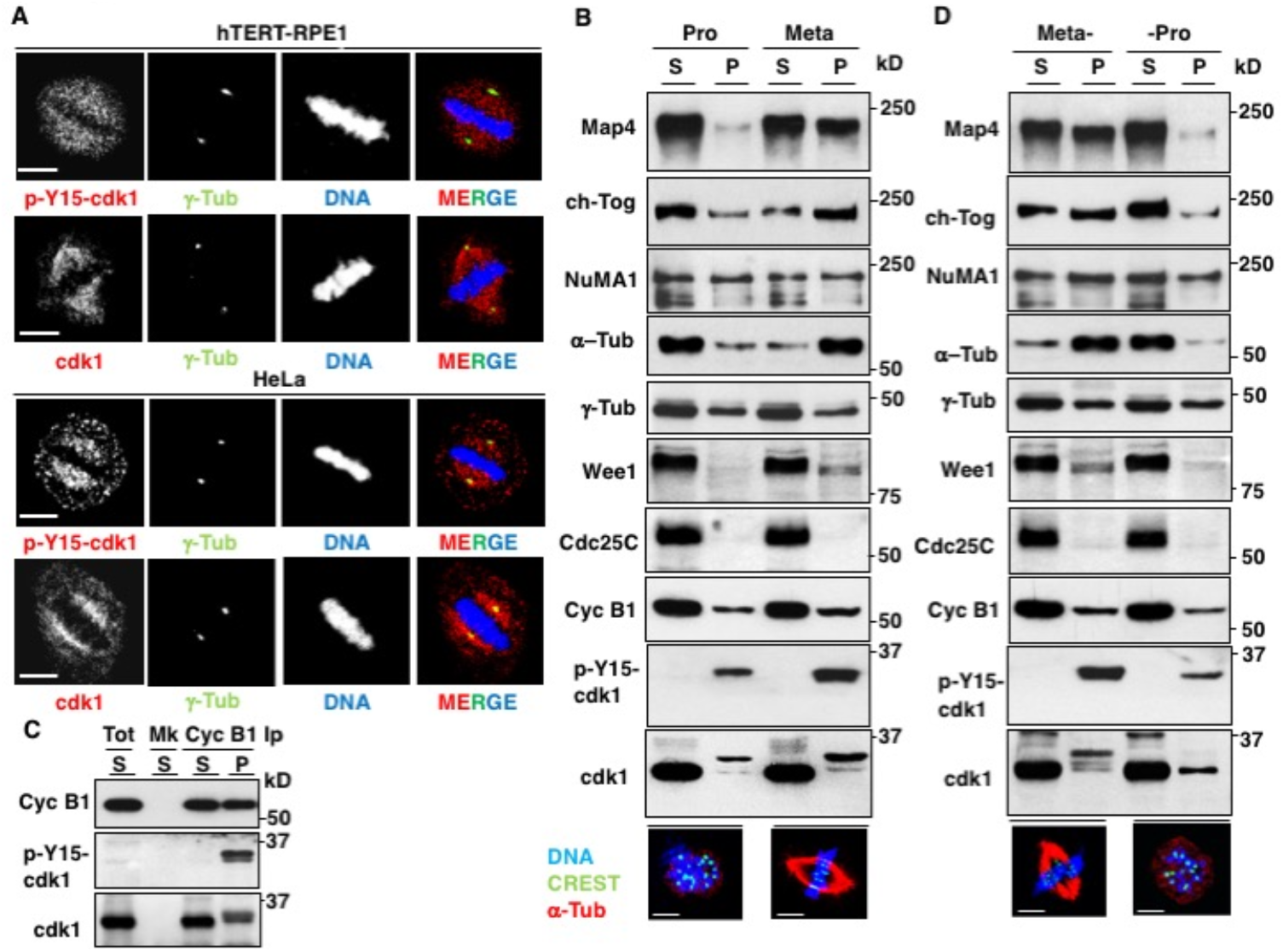
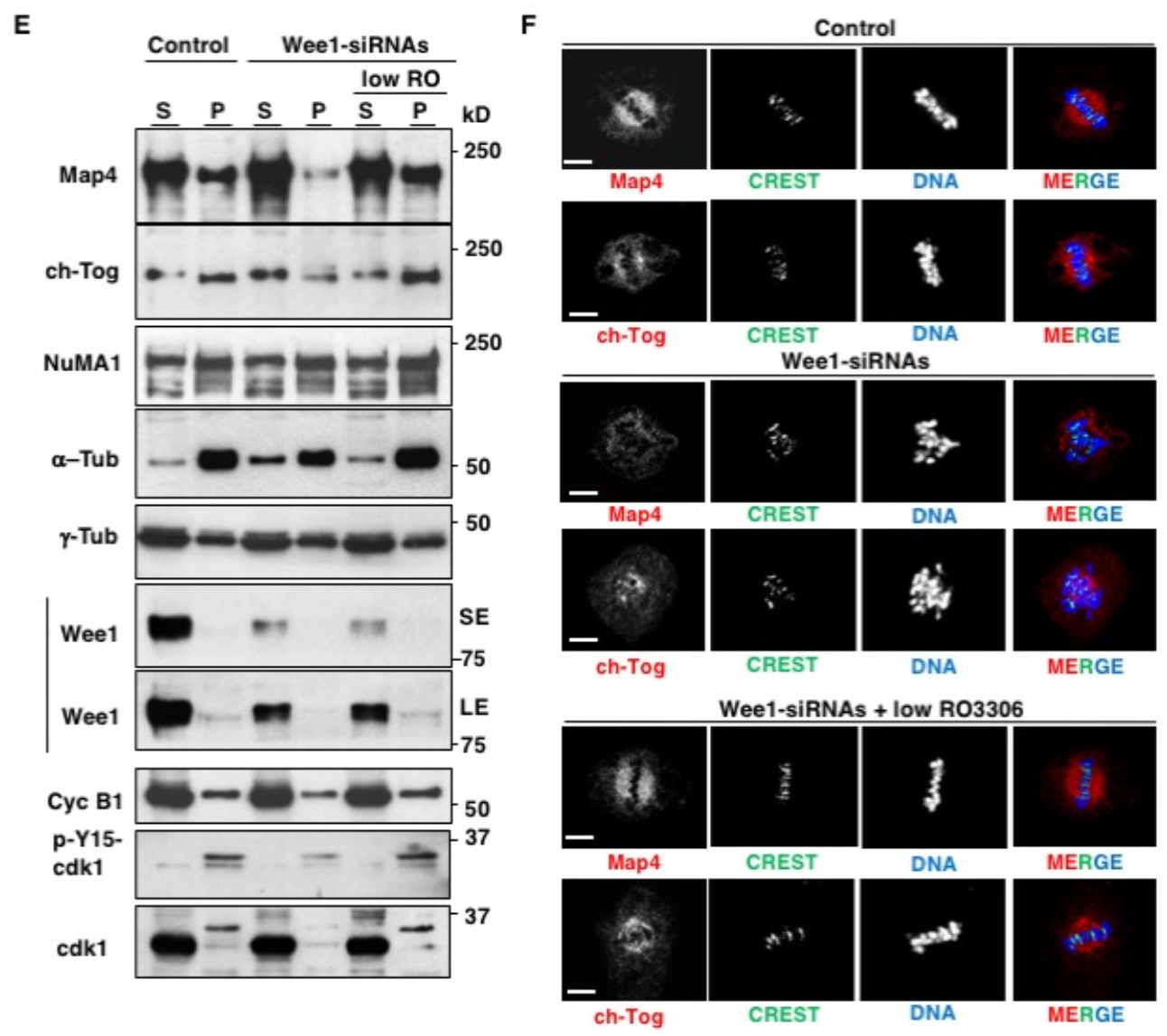
i-Cdk1 localizes at spindle structures and is required for spindle assembly. (**A**) Growing hTERT-RPE1 and HeLa cells were treated for 20 min with MG-132, fixed and stained for indicated antigens (g-Tub; γ-tubulin). (**B**) Prometaphase-arrested hTERT-RPE1 cells were released in M/C medium for 60 min and either nocodazole was added back at time 0 of incubation (Pro), or just vehicle (Meta). S and P fractions were probed for indicated antigens. Bottom: IFs of representative cells probed for indicated antigens. (**C**) Cyclin B1 Ips from Meta cell S and P fractions probed for indicated antigens. (**D**) Meta cells obtained as described in (B) were further treated with vehicle (Meta-) or nocodazole (-Pro) from 60 min to 80 min of incubation. S and P fractions were probed for indicated antigens. (**E**, **F**) hTERT-RPE1 cells were treated with non-targeting-siRNAs (Control) and Wee1-siRNAs, prometaphase-arrested and released in M/C medium for 80 min incubation as described in Fig. 1b and a portion of Wee1-siRNA-treated cells received low-RO at 60 min post release. (**E**) S and P fractions probed for indicated antigens; (**F**) Cells were fixed and stained for indicated antigens. Scale bar: 5 μm.

To ask whether a cause/effect relationship existed between i-Cdk1 and the control of Map4 and ch-Tog dephosphorylation and distribution, prometaphase-arrested control and Wee1-siRNAs were released into M/C medium as described in Fig. 1b. Cells were further fractionated and the S and P fractions analyzed (fig. 2E). Control cells showed a similar distribution as in Meta cells described in Figs. 2b, d. In Wee1-siRNA-treated cells the distribution of γ-tubulin and NuMA1 between the S and P fractions were similar to control cells, however, Map4, ch-Tog and even part of α-tubulin were substantially less present in the P fraction relatively to control cells, a distribution pattern similar to that of Pro cells described in figs 2A, D (fig. 2E). Addition of low-RO to Wee1-siRNA-treated cells reversed protein distribution relocating Map4, ch-Tog and α-tubulin from S to P fraction similarly to control cells (fig. 2E). As expected, i-Cdk1 was reduced in Wee1-dowregulated cell P fraction compared to controls, however, addition of low-RO induced accumulation of part of residual Wee1, that escaped siRNA-mediated downregulation, in P fraction and restored i-Cdk1 content (fig. 2E; note Wee1 in short, SE, and long exposures, LE). No increase of p-Y15-cdk1 signal was detected in S fraction upon low-RO addition, indicating that small fluctuation of Cdk1 activity are indeed buffered in the cytosol by strong hysteresis and bistability of the Cdk1 autoactivatory loops, while nucleating spindle microtubules appeared insulated from them (fig. 2E) (*8, 20, 21*). Localized inhibition of endogenous Cdk1 by low-RO treatment, could possibly explain spindle assembly restoration in cdk1AF-expressing cells (fig. S2). Immunofluorescence (IF) analysis confirmed that the two MAPs poorly localised to defectively structured spindles in Wee1-downregulated cells in a low-RO-reversible manner (fig. 2F). Similar mitotic defects and changes in protein distribution were induced by the chemical Wee1 inhibitor AZD1775 upon release from prometaphase-arrest, also in this case corrected by low-RO (fig. S4B). Together, these data indicate that i-Cdk1 is required for proper mitotic spindle assembly and spindle microtubule association of Map4 and ch-Tog.

Microtubule growth- and stability-promoting properties of MAPs like ch-Tog and Map4 are antagonized by direct Cdk1 phosphorylation (*2, 3, 4, 5, 6*). We were intrigued by the fact that changes in migration on SDS/PAGE suggested that these MAPs were hypophosphorylated when associated with spindle microtubules (fig. 2B, D). We developed an anti-phosphospecific antibody recognizing phosphorylated Map4-serine 787 (p-S787-Map4), a target site for inhibition by Cdk1 *in vivo* (*22*). Indeed, when the fusion proteins tGFP-Map4-wild type (WT) and tGFP-Map4-S787A (S787A), in which Map4 serine 787 was mutated into non-phosphorylatable alanine, were expressed and isolated from mitotic cells, the antibody recognized wild type but not the mutant version (fig. S7). Probing Ips of comparable amounts of endogenous Map4 from S and P fractions of Meta cells for p-S787-Map4 showed that the signal was readily detected from S fraction while hardly from P fraction (fig. 3A). In addition, treating P fraction Map4 with active Cdk1, *in vitro*, restored strong p-S787-Map4 signal and slower Map4 migration (fig. 3B). These data indicate that Map4 is dephosphorylated at Cdk1-dependent sites when bound to microtubules. Map4 and ch-Tog have been shown to physically interact with Cdk1, mediating association of the kinase with microtubules (*6, 23*). Our data from Wee1 downregulation experiments suggested that ch-Tog and Map4 distribution in the P fraction depended on i-Cdk1. Indeed, Ips of comparable amounts of Map4 or ch-Tog from the S and P fraction of Meta cells showed that the MAPs selectively interacted with Wee1 and i-Cdk1 in P fraction (figs. 3C, D). Together, these observations suggest that the interaction of microtubule-associated MAPs with i-Cdk1 was instrumental for their dephosphorylation (fig. 3A, C, D). In mitosis, major protein phosphatases like PP1 and PP2A are directly or indirectly inhibited by Cdk1 (*1*). PP1 catalytic subunit is directly inhibited by Cdk1 phosphorylation, in particular the alpha isoform, PP1α, at threonine 320 (p-T-320-PP1α) and phosphorylation in analogous sites inhibits other PP1 isoforms (*24*). PP1 can, however, rapidly autoactivate by dephosphorylating itself, provided no Cdk1 activity in the proximity (*24*). In addition, PP1 often interacts with its substrates through specific domains (*25*). Thus, first we asked whether PP1 interacted with Map4 and ch-Tog and found that the MAPs interacted with PP1α only in the P fraction (figs. 3C, D). Because of the presence of i-Cdk1, we hypothesised that PP1α in P fraction was dephosphorylated at the Cdk1 inhibitory site. Indeed, when comparable amounts of PP1α were analyzed from S and P fractions of Meta cells, the p-T-320-PP1 signal was substantially detectable in S fraction and barely in P fraction (fig. 3E). In addition, probing comparable amount of PP1 a from total S fraction and from Map4 and ch-Tog Ips showed that MAP-associated PP1α was dephosphorylated at the inhibitory Cdk1-dependent site (fig. 3F). Moreover, active PP1α was able to dephosphorylate Map4-S787 *in vitro* (fig. 3G). Together, these data suggest that i-Cdk1 coopts active PP1 to reverse inhibitory phosphorylation of Map4 and ch-Tog, and possibly other MAPs, for spindle microtubule growth. When prometaphase-arrested control-siRNA- and Wee1-siRNA-treated cells were released into M/C medium for 80 min, PP1α was readily detectable in Map4 and ch-Tog Ips from total extracts of control but not of Wee1-siRNA-treated cells (fig. 3H, lane 6). Addition of low-RO to Wee1-siRNA-treated cells, that restored i-Cdk1 and spindle assembly (see fig. 2E), also restored efficient PP1α binding to Map4 and ch-Tog (fig. 3H, lane 9). Thus, i-Cdk1 allowed Map4 and ch-Tog interaction with active PP1 to promote their dephosphorylation. The Map4 sequence KDVRW (from aa 318 to 322) matched a rather conserved PP1 binding motif (R/KxVxF/W; where x is any aa) (*25*). We mutated it into KDARA in tGFP-Map4-WT expression vector to produce tGFP-Map4-PP1-KO, a mutant impaired in PP1 binding. Indeed, when tGFP-Map4-WT and tGFP-Map4-PP1-KO were expressed at comparable levels and isolated from Meta cells, PP1a substantially bound tGFP-Map4-WT but not tGFP-Map4-PP1-KO (figs. 3J, K). When the distribution in S and P fractions of tGFP-Map4-WT- and tGFP-Map4-PP1-KO-transfected Meta cells was analyzed, tGFP-Map4-WT was readily detectable in the P fraction while tGFP-Map4-PP1-KO remained substantially in S fraction (fig. 3I). We concluded that i-Cdk1 is required for spindle localized interaction of microtubule-stabilizing MAPs with active PP1 and for PP1-dependent reversal of their inhibitory phosphorylations, operated by Cdk1 and other mitotic kinases, in order to locally restore their microtubule growth-promoting properties.

**Fig. 3.**
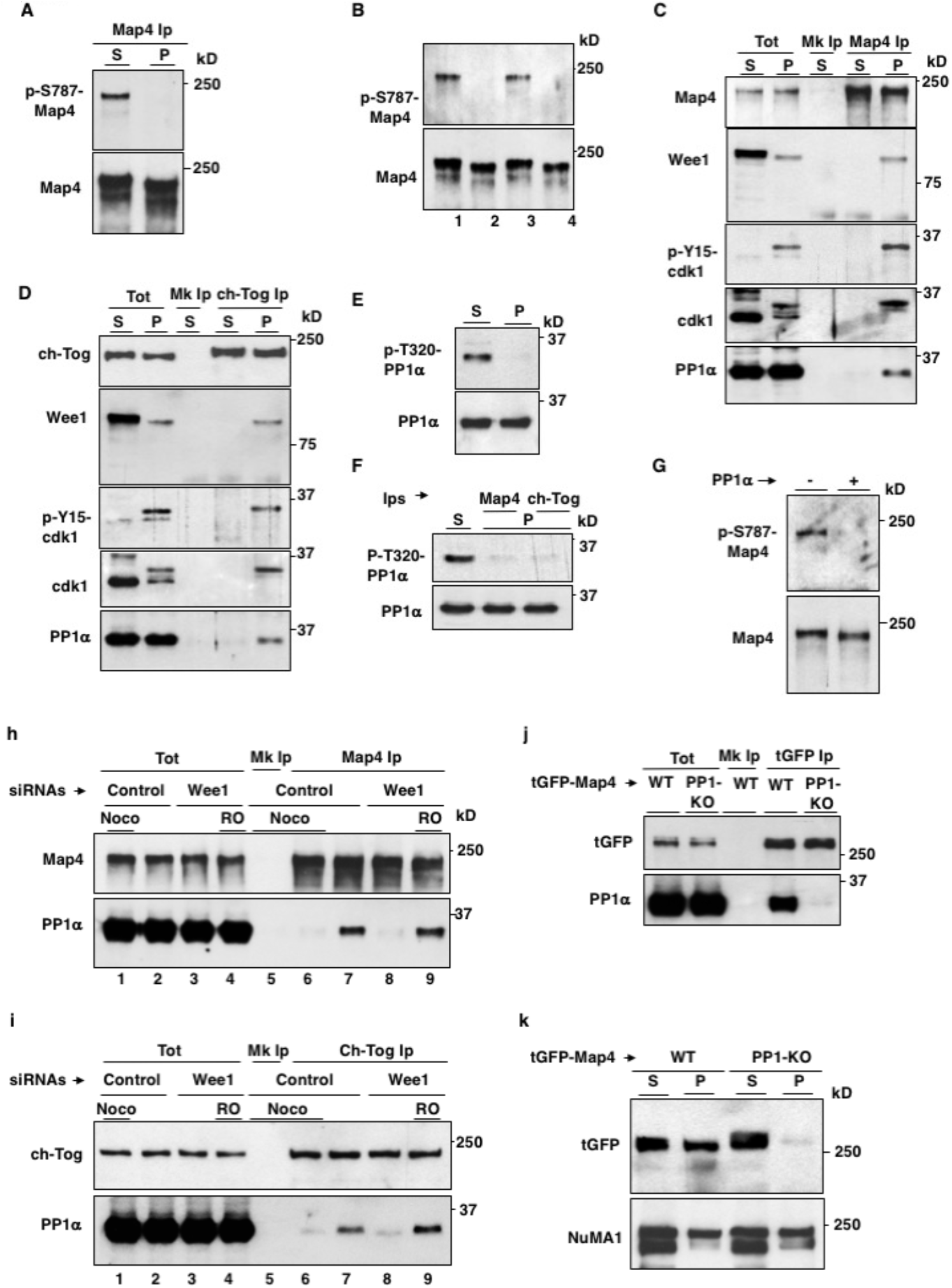
i-Cdk1 assists spindle-localized MAP interaction with PP1 and dephosphorylation. From Meta hTERT-RPE1 cells: (**A**) Map4 Ips from S and P fractions were probed for indicated antigens. (**B**) Map4 Ips were incubated in kinase reaction buffer at 37° C for 15 min: lane 1, from S fraction; lanes 2, 3 and 4, from P fraction – or + active Cdk1 or + active Cdk1 + RO-3306, respectively. After incubation, Ips were probed for indicated antigens. (**C**) Map4 Ips and (**D**) ch-Tog Ips from S and P fractions were probed for indicated antigens. **e**, S and P fractions were probed for indicated antigens. (**F**) total S fraction and Map4 and ch-Tog Ips from P fraction were probed for indicated antigens. (**G**) Map4 Ips from S fraction were incubated in phosphatase reaction buffer at 37° C for 15 min in - or + active PP1α, then probed for indicated antigens. (**H**) and (**I**) Non-targeting-siRNA-(Control) and Wee1-siRNA-treated (Wee1) hTERT-RPE1 cells released from prometaphase-arrest into M/C medium for 80 min incubation. A sample of control cells received nocodazole at time 0 (Noco) and a sample of Wee1-siRNAs cells received low-RO at 60 min of incubation (RO). Cells were lysed in high salt buffer and (**H**) Map4 or (**I**) ch-Tog Ips were probed for indicated antigens (lanes 1-4, total lysate; Mk Ip, mock Ips from Noco cell lysates). (**J**, **K**) hTERT-RPE1 cells were transfected with tGFP-Map4-wild type (WT) and tGFP-Map4-PP1-KO expression vectors, arrested at metaphase and either (J) lysed in high salt buffer, tGFP Ips were probed for indicated antigens or (K) S and P fraction isolated and probed for indicated antigens.

Probing the cyclin B1 Ips form S and P fractions of Meta cells, shown in fig. 2C, for γ-tubulin revealed that it interacted with Cdk1 in P fraction (fig. 4A). Conversely, probing γ-tubulin Ips from S and P fraction of Pro cells showed that it selectively interacted with i-Cdk1 and Wee1, in P fraction (fig. 4B). In Meta cells P fraction, i-Cdk1 and Wee1 increased relatively to the P fraction of Pro cells, suggesting a mutual relationship between microtubule polymerization and i-Cdk1 formation (see figs. 2B, D and 3C, D). This may reflect further Wee1 recruitment and i-Cdk1 formation at intra-spindle microtubule nucleation points, in a γ-tubulin-dependent manner, but also further i-Cdk1 formation by growing spindle microtubules, in a mutual fashion, as suggested by the interaction of i-Cdk1 with Map4 in Meta cell P fraction (see fig. 3C) (*17, 18*). When the repartition of the Cdk1 inhibitory pathway players between S and P fractions in Pro and Meta cells was probed on blots where the P fraction was enriched 10 folds over the S fraction, it appeared more evident that the increase of Wee1 in P fraction of Meta cells relatively to Pro cells was microtubule-dependent, Myt1 also increased albeit to a lesser extent (fig. 4C). No changes in distribution between fractions in Meta and Pro cells were, instead, detected for Cdc25 phosphatase family members (fig. 4C). Thus, polymerizing spindle microtubules appeared to bind Wee1 but not with Cdc25, further promoting i-Cdk1 insulated from the cytoplasmic Cdk1 autoactivatory loops (fig. 4C) (*8, 20, 21*). To corroborate the hypothesis that nucleating microtubules compartmentalized the Cdk1 control fostering i-Cdk1 formation, we used concentrated cytoplasmic cell extracts from prometaphase-arrested HeLa that were incubated with nocodazole or with the microtubule stabilizer taxol for 20 min at 23°C, the pellets were then eluted and analyzed (*25*). The microtubular pellets aggregated with Wee1 and i-Cdk1 but not with Cdc25 family members (fig. 4d). The presence of Wee1 on spindles in cells was also confirmed by IF (fig. 4E). Thus, in mitosis, positive feedback loops keep Cdk1 active in the cytoplasm to clear peripheral cytoplasmic microtubules by phosphorylating MAPs for safe chromosome movements and possibly to destabilize spindle microtubules that do not, or weakly, bind chromosomes (fig. 4F). At the same time, centrosomes, and perhaps intra-spindle γ-tubulin nucleating centers, along with their nucleating microtubules foster localized i-Cdk1 for PP1-dependent reversal of inhibitory MAP phosphorylation and spindle microtubule growth (fig. 4f) (*3, 17, 18, 27*). Compartmentalized control of Cdk1 activity might also be contributed by other mechanisms in addition to inhibitory phosphorylation of Cdk1 and relevant to terminate the SAC-sustaining action of Cdk1 upon stable microtubule-kinetochore attachments (*15, 28, 29*).

**Fig. 4.**
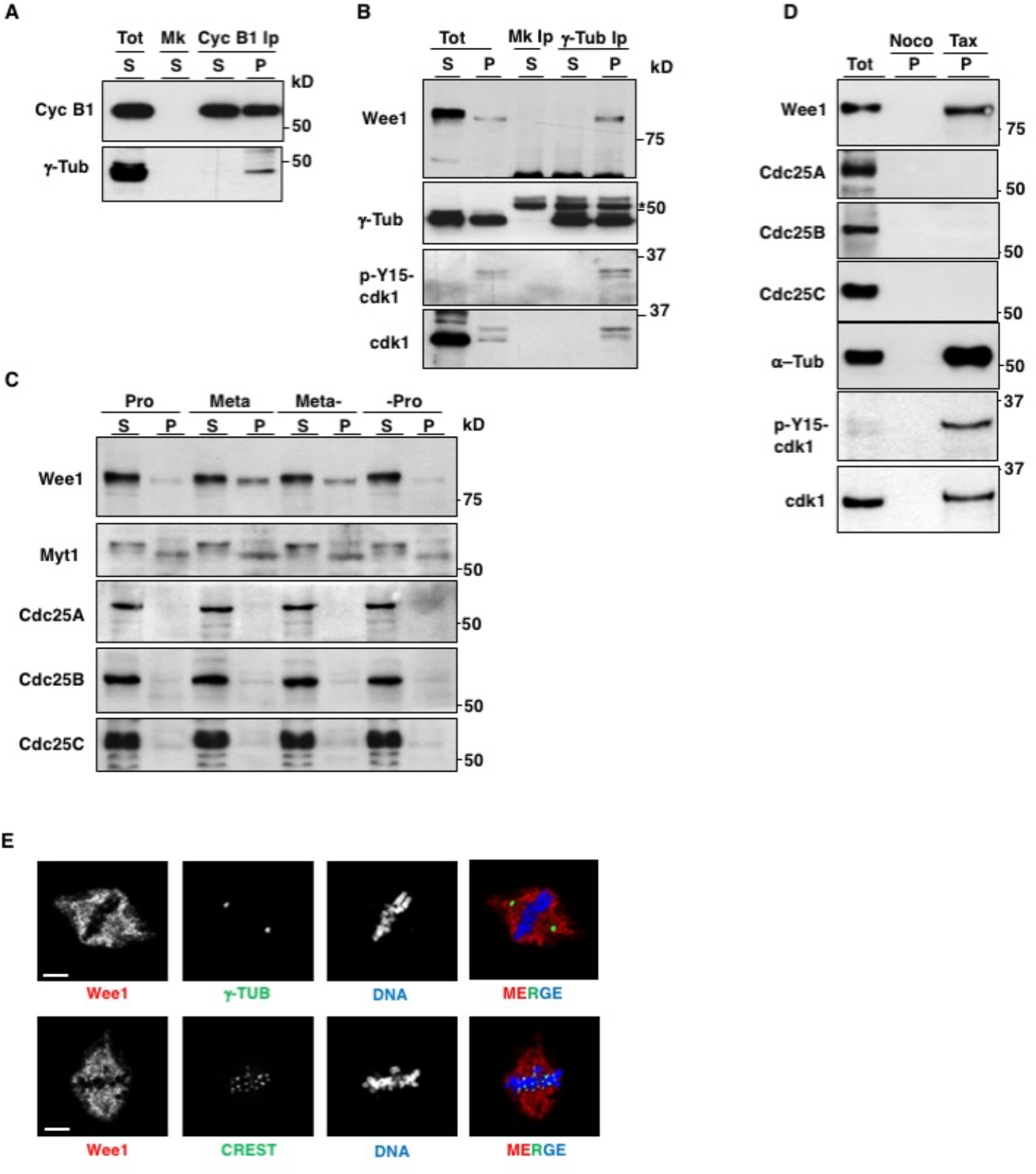
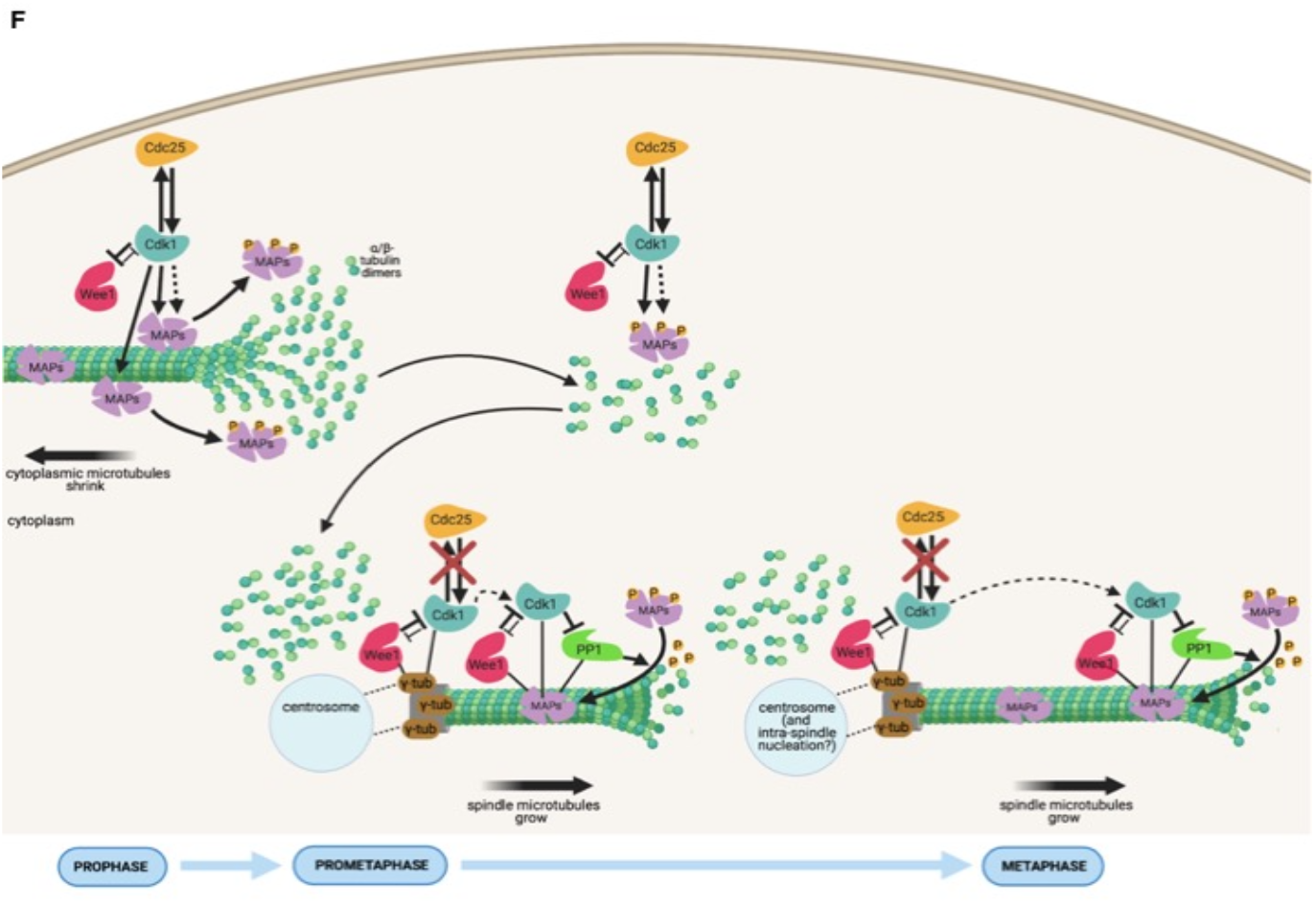
Nucleating microtubules bind and foster i-Cdk1. (**A**) Cyclin B1 Ips described in fig. 2C were probed for γ-tubulin. (**B**) γ-tubulin Ips from S and P fraction of Pro hTERT-RPE1 cells were probed for indicated antigens (P fraction enriched 10 folds over S fraction). (**C**) S and P fraction from Pro, Meta, Meta- and -Pro, as described in Fig. 2b, d, were probed for indicated antigens on blots in which the P fraction was enriched 10 folds over S fraction. (**D**) Eluates from pellets of mitotic HeLa cell extracts (P) incubated with nocodazole (Noco) or taxol (Tax) were probed for indicated antigens; Tot: 1/20 of total extract. **E**, growing hTERT-RPE1 cells were treated for 20 min with MG-132, fixed and stained for indicated antigens. Scale bar: 5 μm. **(F**) a synoptic scheme depicting Cdk1 activity control in mitosis. At prophase, Cdk1 activity, sustained by positive feedback loops in the cytoplasm, destabilizes cytoplasmic microtubules by directly (continuous arrow) or indirectly (dotted arrow) phosphorylating MAPs. Centrosomes, however, maintain/promote inhibitory phosphorylation of a small Cdk1 fraction by selectively interacting with Wee1 and excluding Cdc25. From prometaphase to metaphase, further interaction of Wee1, inhibited Cdk1 and active PP-1 with microtubule-stabilizing MAPs expands from centrosomes, and possibly from intra-spindle γ-tubulin-dependent nucleating centers, to promote reversal of inhibitory MAP phosphorylation and spindle microtubule growth.

## Acknowledgements

The authors acknowledge V. E. Avvedimento for suggestions, R. Visconti and A. Romano for technical advice. D.G. acknowledges AIRC, Associazione Italiana per la Ricerca sul Cancro, for support (IG grant 2017; Id. 19851 to D.G.).

## Author contributions

A.F.S. and D.G. designed and performed experiments and analyzed data. C.P. and F.F. performed immunoblot and immunofluorescence experiments and analyzed data. D. G. wrote the manuscript. A.F.S. revised the manuscript. All authors read and approved the final manuscript.

## Supplementary Materials

### Materials and Methods

#### Cell lines and cell culture

HeLa cells were grown in Roswell Park Memorial Institute Medium (RPMI-1640; Sigma-Aldrich), hTERT-RPE1 cells were grown in Dulbecco’s Modified Eagle Medium: Nutrient Mixture F-12 (DMEM/F12; Gibco, Thermo Fisher Scientific), both supplemented with 10% fetal bovine serum (FBS; Gibco), 1% GlutaMAX-supplement (Gibco), 1% penicillin/streptomycin (Euroclone), at 37° C with 5% CO_2_.

#### Cell treatments

Transfections were performed using linear polyethylenimine (PEI; Polysciences, Inc) or MegaTran 2.0 Transfection Reagent (OriGene Technologies, Inc.). The expression plasmids for turbo-GFP-Map4-wild type (tGFP-Map4-WT), turbo-GFP-Map4 rendered non-phosphorylatable at serine 787 (tGFP-Map4-S787A) and turbo-GFP-Map4 defective in PP1 binding Map4 (tGFP-Map4-PP1-KO), in which the Map4 peptide KDVRW (from aa 318 to 32; human Map4 isoform 1) was mutated into KDARA, were generated and purchased from OriGene Technologies Inc. Cdk1WT and cdk1AF expression plasmids were purchased from Addgene. 3XFlag-Wee1 expression construct was obtained as described^13^. SiRNAs targeting the 3’-UTR of human WEE1 (5’-GGGCUUU AUUACAGACAUAUU-3’, 5’-GUACAUAGCUGUUUGAAAUUU-3’ and 5’-UGUAAACUUGUAGCAUUAUU-3’) were purchased from Dharmacon Inc. For efficient knockdown cells were transfected with 25 nM of siRNAs duplex using DharmaFECT 1 siRNA Transfection Reagent (Dharmacon Inc.). For siRNA treatment and complementation experiments in synchronized cells, cells were mock-or 3XFlag-Wee1 expression construct-transfected 24 hours prior to treatment with non-targeting or specific siRNAs. For synchronization of siRNA-treated cells, 6 hours after siRNA-treatment, cells were incubated with RO-3306 at 9 μM (high-RO; Calbiochem) for further 16 hours of incubation, to have G2-arrsted cells, or with nocodazole (Abcam) at 1 μg/ml and 100 ng/ml for further 16 or 14 hours of incubation for hTERT-RPE1 and HeLa cells, respectively, to have prometaphase-arrested cells (Pro cells). To obtain metaphase-arrested cells (Meta cells), Pro cells were collected and washed twice with fresh medium and twice with phosphate buffer saline (PBS; Euroclone) solution before plating into fresh medium containing MG-132 (20 μM; Calbiochem) and cycloheximide (CHX; 60 μg/ml; Santa Cruz Biotechnology, Inc) for further 60 min incubation. Meta- and -Pro cells were obtained by adding either DMSO or nocodazole (1 μg/ml) to Meta cells at 60 min and prolonging incubation for further 20 min. Where indicated AZD1775 (Selleckchem) was added at 1 μM. Where indicated, partial reversal of Cdk1 activity was obtained by adding RO-3306 at 0.5 μM (low-RO).

#### Cell fractionation and extracts

To isolate spindle-bound protein we adapted a previously described method^15^. All the steps, except the final elution, were performed at room temperature (rt) to preserve microtubules. Cells were harvested by centrifugation and washed once with PBS containing taxol (1 μM; Sigma-Aldrich). 5 x 10^6^ cells were resuspended in 800 μl of fractionation lysis buffer (FLB; 40 mM β-glycerophosphate, 15 mM MgCl2, 20 mM EGTA, 0,25% Igepal, 20 mM Hepes, 5 μM taxol, 6 μg/ml latrunculin B; Sigma-Aldrich) supplemented with 300 μg/ml RNase A and 120 U/ml DNase I (Roche). After careful resuspension, samples were incubated in thermomixer (Eppendorf thermomixer comfort) at 34 °C for 20 min with constant shaking at 1,200 rpm. Lysates were spun at 6,800 rpm (Eppendorf centrifuge 5425) for 2 min: the supernatant fractions were collected and transferred into new tubes and supplemented with 250 mM NaCl (S fraction), the pellet fractions were resuspended in FLB + 300 μg/ml RNase A and 120 U/ml DNase I and again incubated in thermomixer at 34 °C for 20 min with constant shaking at 1,200 rpm. Lysates were collected again by centrifugation and the pellets were washed twice with washing buffer (WB; 5 mM Hepes, 5 μM taxol). Finally, the pellet fractions were resuspended in 200 μl of FLB minus taxol and supplemented with 250 mM NaCl (P fraction). Both supernatant and pellet fractions were incubated on ice, to help destabilize microtubules, for 30 min and then spun at 3,200 rpm at 4 °C for 10 min (Eppendorf centrifuge 5424 R) and the supernatants of S and P fraction further analyzed by Ibs or Ips. To induce microtubule polymerization and analyze microtubule-bound proteins in mitotic cell extracts, HeLa cell extracts were obtained as previously described^24^. Cell extracts were supplemented with 1 mM GTP (Sigma-Aldrich), 50 μl aliquots were treated with nocodazole (1 μg/ml; Abcam), as control, or taxol (5 μM; Sigma-Aldrich) for 20 min at rt, diluted 10 times with FLB and spun at 6,800 rpm (Eppendorf centrifuge 5425) for 2 min. The pellets were washed once more in FLB and finally resuspended in 30 μl FLB supplemented with 250 mM NaCl, incubated on ice for 30 min and then spun 13,200 rpm at 4 °C for 10 min (Eppendorf centrifuge 5424 R). Supernatants were analyzed by Ibs; total samples = 5 μl of total mitotic extract.

#### Antibodies

Antibodies used for immunoblotting according to standard protocols: mouse anti-cdc2 (1:500; BD Biosciences), rabbit anti-phospho-tyrosine-15-cdc2 (P-Y 15-cdc2; 1:1000; Boster Bio), rabbit anti-phospho-threonine-14-cdc2 (P-T14-cdc2; 1:1000; Cell Signaling Technology Inc.; CST), rabbit anti-Wee1 (1:1000; CST), rabbit anti-Myt1 (1:1000; CST), rabbit anti-Cdc25C (1:1000; CST), rabbit anti-phospho-threonine-320-PP1α (P-T320-PP1; 1:1000; CST), rabbit anti-cyclin B1 (1:2000; Bethyl Laboratories, Inc.), mouse anti-g-tubulin (1:2000; Sigma-Aldrich), mouse anti-α-tubulin (1:2000; Sigma-Aldrich), mouse anti-PP1g (1:500; Santa Cruz Biotechnology, Inc.), rabbit anti-Map4 (1:2000; Bethyl Laboratories Inc.), rabbit anti-NuMA (1:1000; Novus Biologicals), rabbit anti-ch-Tog (1:1000; Abcam), mouse anti-turboGFP (1:2000; OriGene Technologies). Rabbit polyclonal antibody against phosphorylate serine 787 of human Map4 (P-S787-Map4) was raised using C-DAKAPEKRA[Sp]PSKPA-coNH2 peptide as immunogen; peptide synthesis, rabbit inoculation, serum production and cross-affinity antibody purification were carried out by CovalAb S.A.S. Sheep anti-mouse IgG peroxidase-(HRP) linked and donkey anti-rabbit IgG HRP-linked whole antibodies (1:2000; GE Healthcare). Antibodies used for IF: rabbit anti-phospho-tyrosine-15-cdc2 (P-Y 15-cdc2; 1:50; CST), anti-cdc2 (1:100; BD Biosciences), mouse anti-g-tubulin (1:1000; Sigma-Aldrich), mouse anti-α-tubulin (1:1000; Sigma-Aldrich), rabbit anti-g-tubulin (1:1000; Sigma-Aldrich), human anti-centromere (CREST; 1:50; Antibodies Incorporated), rabbit anti-Map4 (1:1000; Abcam), rabbit anti-ch-Tog (1:400; Proteintech Group, Inc.), rabbit anti-Wee1 (1:100; CST), goat anti-human IgG rhodamine conjugated (4 μg/ml; Santa Cruz Biotechnology, Inc.), rabbit anti-human IgG FITC conjugated (4 μg/mL; Dako Agilent), Alexa Fluor 594 donkey anti-rabbit IgG (2 μg/mL; Invitrogen), Alexa Fluor 594 donkey anti-mouse IgG (4 μg/ml; Invitrogen), Alexa Fluor 488 donkey anti-rabbit IgG (2 μg/ml; Invitrogen), Alexa Fluor 488 goat anti-mouse IgG (4 μg/ml; Invitrogen). Antibodies used for Ips: rabbit anti-Map4 (2μg/mL; Bethyl Laboratories Inc.), rabbit anti-chTOG (2μg/mL; Abcam), mouse anti-g-tubulin (2μg/mL; Sigma-Aldrich). Antibody-conjugated beads used for Ips: mouse anti-cyclin B1 agarose (Santa Cruz Biotechnology), mouse anti-turboGFP agarose (OriGene), protein A/G PLUS-agarose (Santa Cruz Biotechnology), mouse anti-g-tubulin agarose conjugate (Santa Cruz Biotechnology).

### Immunoblotting and immunoprecipitation

Immunoblots (Ib) were performed as described^26^. For immunoprecipitations (Ips) from S fraction and and P fraction eluates, samples were diluted 6 folds in lysis buffer (LB; 80 mM β-glycerophosphate, 15 mM MgCl2, 20 mM EGTA, 20 mM Hepes pH 7.4, 100 mM NaCl, 0,1 % Igepal; Sigma-Aldrich) supplemented with 1 tablet of phosphatase inhibitor cocktail (PhosSTOP; Roche) and incubated with antibody and protein A/G PLUS-agarose beads or with agarose bead-conjugated antibody overnight at 4 °C in constant rotation. Beads were washed twice in LB and beads-bound proteins were eluted by boiling in SDS loading buffer. For total cell lysates, 5 x 10^6^ cells were lysed in 180 μl high salt lysis buffer (HSB; 16 mM β-glycerophosphate, 3 mM MgCl2, 4 mM EGTA, 0,5 mM DTT, 20 mM Hepes, 0,2 % Igepal, 250 mM NaCl; Sigma-Aldrich) supplemented with 1 tablet of phosphatase inhibitor cocktail (PhosSTOP; Roche) to preserve phosphorylation. After 30 min incubation on ice, lysates were spun at 13,200 rpm for 10 min at 4° C (Eppendorf centrifuge 5424 R). Cleared lysates were incubated with antibody and protein A/G PLUS-agarose beads or with agarose bead-conjugated antibody overnight at 4 °C in constant rotation. Beads were washed twice in LB and beads-bound proteins were eluted by boiling in SDS loading buffer. For kinase reactions, after the two LB washes, Ips were washed once in kinase reaction buffer (KRB; 80 mM β-glycerophosphate, 15 mM MgCl2, 20 mM EGTA, 20 mM Hepes pH 7.4, 100 μM ATP; Sigma-Aldrich) and incubated for 20 min at 37° C in KRB -/+ recombinant active Cdk1 (0.05 mg/ml; Sigma-Aldrich) and + active Cdk1 + RO-3306 (10 μM). For phosphatase reactions, after the two LB washes, Ips were washed once in phosphatase reaction buffer (PRB; 20 mM Hepes pH 7.4, 150 mM NaCl, 1 mM DTT, 1 mM MnCl2; Sigma-Aldrich) and incubated in PRB -/+ 0.01 mg/ml recombinant human PP1α (Novus Biologicals) for 20 min at 30° C.

### Immunofluorescence staining and microscopy

Cells were grown onto poly-D-lysine (0,1 mg/ml; Sigma-Aldrich) coated glass coverslips and treated as described in the text. Before immunofluorescence (IF) procedure initiation, coverslips were washed twice with PBS, then cells were fixed with 4 % paraformaldehyde (Sigma-Aldrich) in PBS (Euroclone) for 10 min at rt and with a methanol (99,9 %; Exacta + Optech) additional step at −20° C for 10 min, the latter except for α-tubulin staining. Cells were permeabilized with 0,25-0,5 % Triton X-100 (Sigma-Aldrich) in PBS for 15-30 min, washed once with PBS and incubated with 3% bovine serum albumin (BSA; Sigma-Aldrich) in PBS for 1 hour at rt. Coverslips were transferred into a humidity chamber and incubated with primary antibodies in 1,5 % (w/v) BSA-PBS solution for 2-12 hour at 4° C. Samples were washed three times and incubated with fluorescently labelled secondary antibodies, dissolved in 1,5 % BSA-PBS solution, for 1 hour at rt. DNA was stained with Hoechst 33258 (1 μg/ml; Invitrogen) by incubation for 10 min. Finally, samples were washed four times and mounted with Mowiol 40-88 (Sigma-Aldrich) on a glass side. Fixed cells were observed using LSM 980 inverted confocal microscope (Zeiss) with a 63X/1,4 oil objective. Representative images were obtained collecting 3 Z-stack series over 42 μm. The acquisitions were deconvoluted and projected into one plane using the LAS-AF software.

### Statistics and reproducibility

For quantitative analyses of immunofluorescence experiments investigators were blinded to sample allocation. Metaphases were visually scored and those with more than three chromosomes falling outside the two internal quarters of the interpolar distance were considered misaligned; around 100 cells were scored in 4 fields of each slide for each experimental condition; each experiment was repeated 3-5 times with similar results. Experiments analyzed by immunoblotting were repeated 2-8 times with similar results.

## Supplementary Figures

**Fig. S1.**
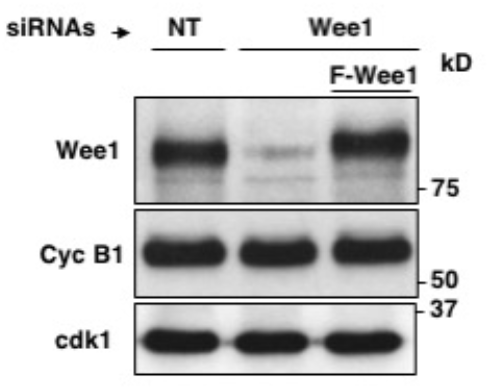
siRNAs treatment of hTERT-RPE1 cells. Lysates of hTERT-RPE1 cells treated with non-targeting siRNAs (NT) and with Wee1-siRNAs (Wee1) as described in fig. 1 were probed for indicated antigens. A portion of Wee1-siRNAs was previously transfected with a siRNAs-resistant, Flag-tagged Wee1 expression vector (F-Wee1).

**Fig. S2.**
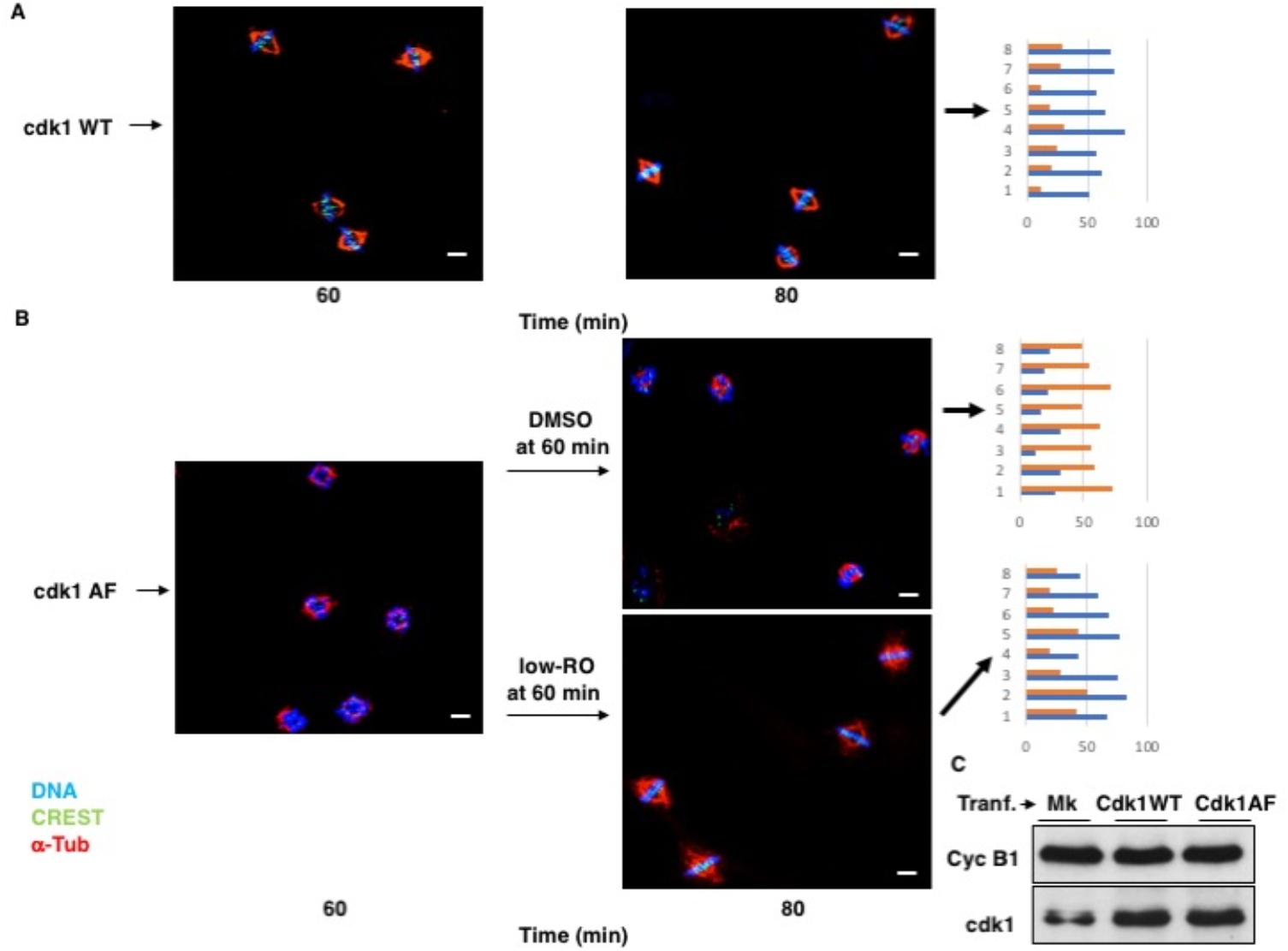
Expression of an inhibitory-phosphorylation-resistant version of Cdk1 impairs spindle assembly. hTERT-RPE1 cells were transfected with (**A**) wild type cdk1 (cdk1WT) expression vector or (**B**) a mutant cdk1 expression vector in which cdk1-threonine 14 and tyrosine 15 are mutated into non-phosphorylatable alanine and phenylalanine, respectively (cdk1AF). 6 hours post transfection, cells were arrested at G2 by addition high-RO and for further 16 hours incubation. Cells were fixed and stained for DNA (blue), α-tubulin (red) and CREST (green) at 60 and 80 min upon release from G2 arrest. A portion of cdk1AF-transfected cells received low-RO at 60 min; as control vehicle was added (DMSO). Graphs: quantitation of cells containing normal aligned (blue bar) or abnormal misaligned (orange bar) spindles at 80 min incubation; shown are data from two independent experiments, in which around 100 cells were scored in 4 independent microscopy slide fields (1-4, experiment 1; 5-8, experiments 2). Scale bar: 5 μm. (**C**) samples of cells treated as in (A) and (B) were lysed prior to release and probed for indicated antigens.

**Fig. S3.**
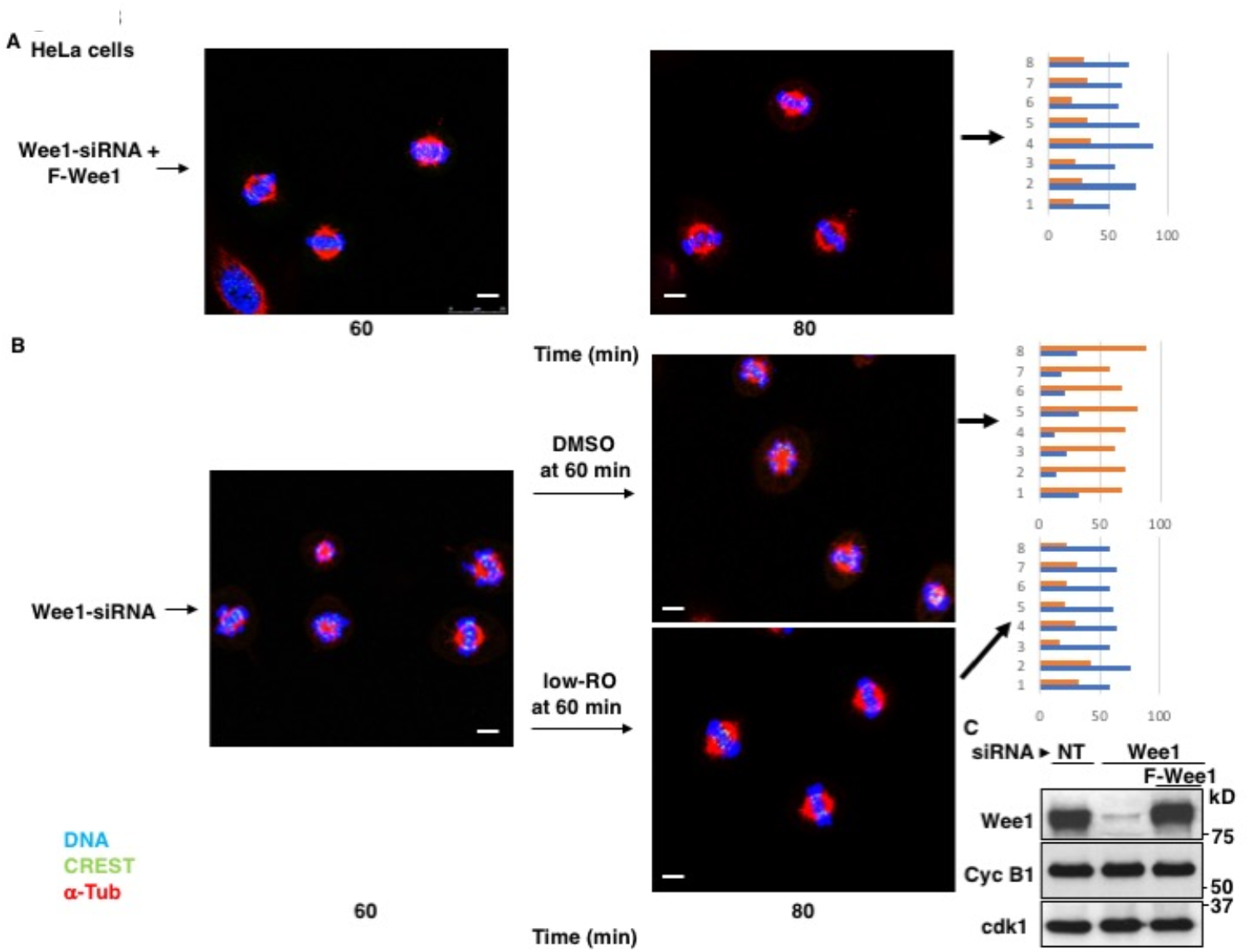
Downregulation of Wee1 impairs spindle assembly in HeLa cells. HeLa cells were treated and arrested at G2 as described for hTERT-RPE1 cells in Fig 1a. Cells were fixed and stained for DNA (blue), α-tubulin (red) and CREST (green) at 60 and 80 min upon release from G2 arrest. (**A**) Wee1-siRNA-treated cells complemented with siRNA-resistant expression vector (Wee1-siRNAs + F-Wee1). (**B**) Wee1-siRNA-treated cells (Wee1-siRNAs). A portion of Wee1-siRNAs cells received low-RO at 60 min; as control vehicle was added (DMSO). Graphs: quantitation of cells containing normal aligned (blue bar) or abnormal misaligned (orange bar) spindles at 80 min incubation; shown are data from two independent experiments, in which around 100 cells were scored in 4 independent microscopy slide fields (1-4, experiment 1; 5-8, experiments 2). Scale bar: 5 μm. **c**, samples of cells treated as in (A) and (B) were lysed prior to release and probed for indicated antigens.

**Fig. S4.**
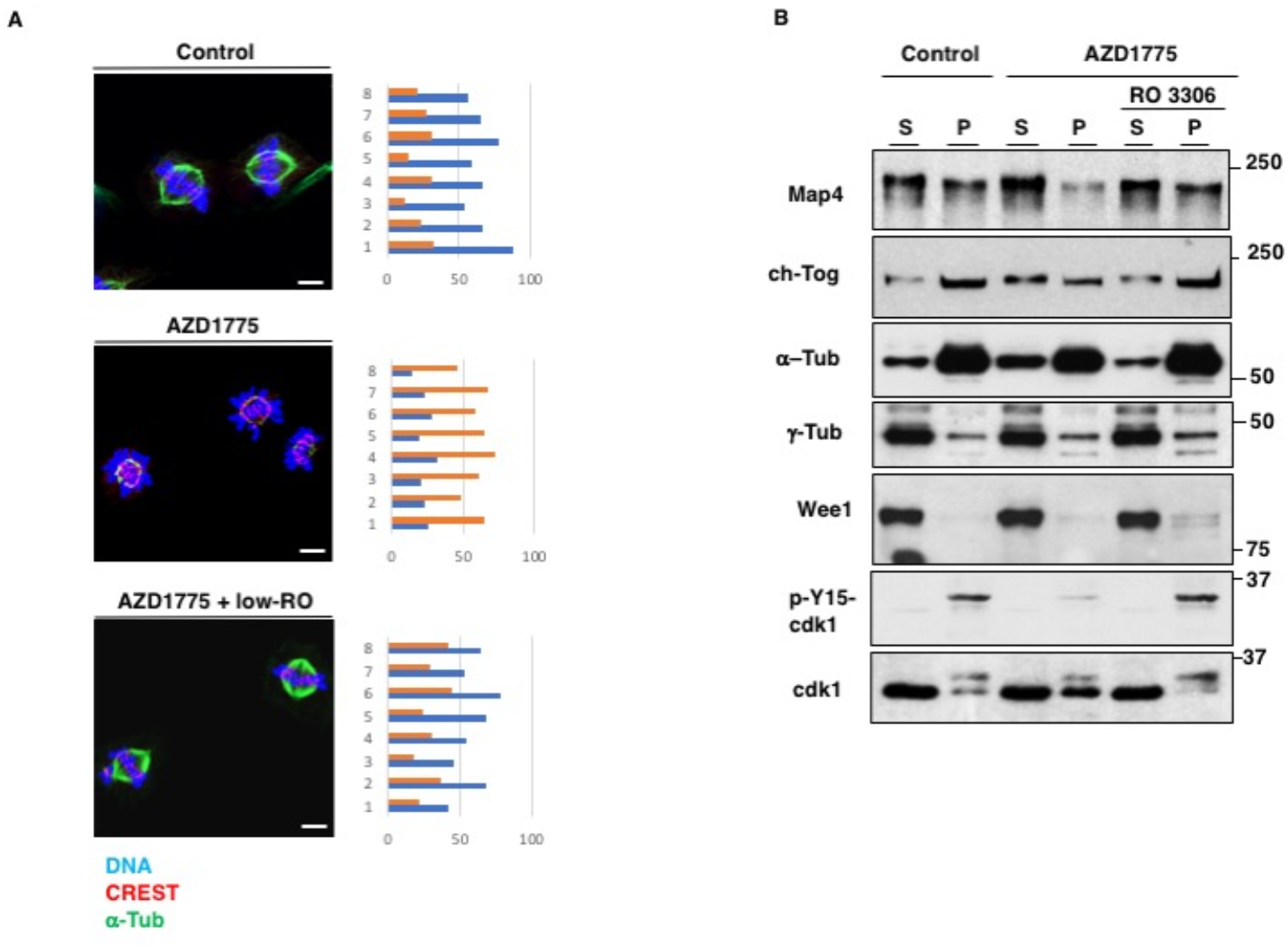
Chemical inhibition of Wee1 impairs spindle assembly. (**A**) hTERT-RPE1 cells were released from a G2 arrest by high-RO treatment into M/C medium and either vehicle (Control) or AZD1775 was added. Cells were fixed after further 80 min of incubation and stained for α-tubulin (green), centromeres (red) and DNA (blue). A sample of the AZD1775-treated cells also received low-RO 60 min post release. Scale bar: 5 μm. (**B**) prometaphase-arrested hTERT-RPE1 were released into M/C medium and either vehicle (Control) or AZD1775 was added. S and P fraction were prepared after further 80 min of incubation and probed for indicated antigens. A sample of the AZD1775-treated cells also received low-RO 60 min post release.

**Fig. S5.**
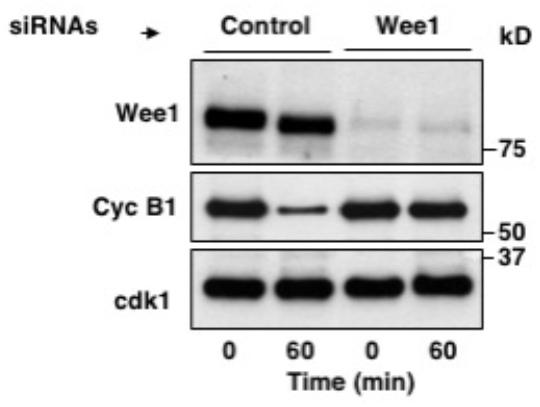
Downregulation of Wee1 delays mitosis exit. Non-targeting-siRNA-(Control) or Wee1-siRNA-treated (Wee1) hTERT-RPE1 cells were arrested at prometaphase and released into fresh medium for 60 min of incubation. Cells were lysed and probed for indicated antigens.

**Fig. S6.**
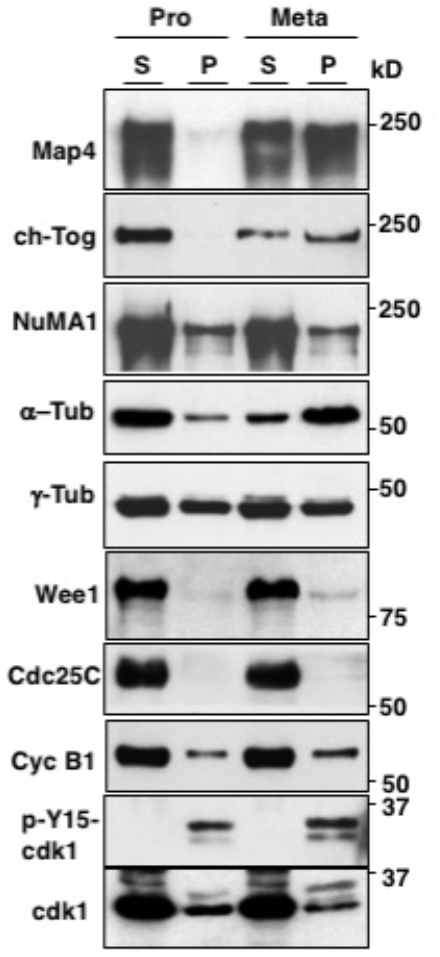
Mitotic HeLa cell fractionation. Prometaphase-arrested HeLa cells were collected and taken immediately (Pro) or released in M/C medium for 60 min (Meta). Fractionated S and P fractions were probed for indicated antigens (P fraction enriched 4 folds over S fraction).

**Fig. S7.**
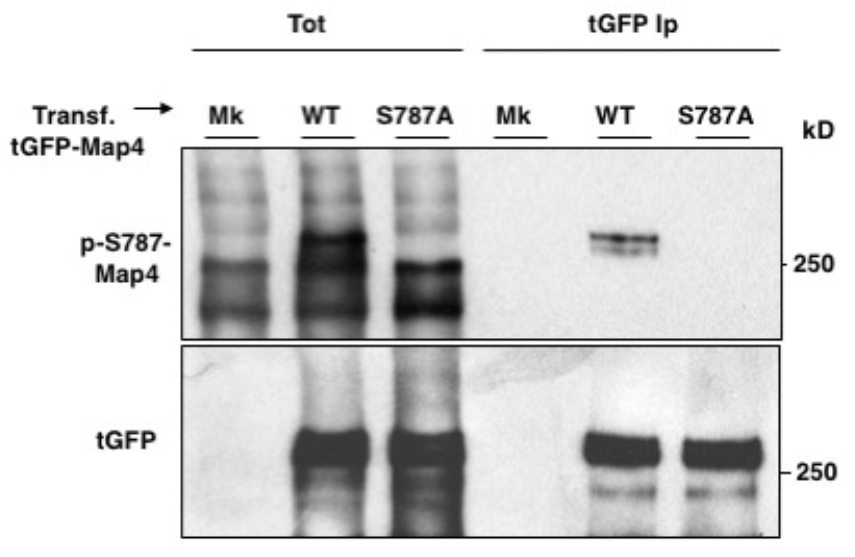
Detection of S787-Map4 phosphorylation. hTERT-RPE1 cells were transfected with expression vectors of tGFP-Map4 wild type (WT) or tGFP-Map4 mutant version in which serine 787 of Map4 was mutated into alanine (S787A) and arrested at prometaphase. Total cell lysates (Tot) and relative anti-tGFP Ips were probed for indicated antigens (Mk: mock-transfected cells).

## References

1. F. Verde, J. C. Labbé, M. Dorée, E. Karsenti, Regulation of microtubule dynamics by cdc2 protein kinase in cell-free extracts of Xenopus eggs. Nature. 343, 233–238 (1990).

2. A. Lieuvin, J. C. Labbé, M. Dorée, D. Job, Intrinsic microtubule stability in interphase cells. J Cell Biol 124, 985–996 (1994).

3. N. Mchedlishvili, H. K. Matthews, A. Corrigan, B. Baum, Two-step interphase microtubule disassembly aids spindle morphogenesis. BMC Biol 16, 14 (2018).

4. Chang, W. et al. Phosphorylation of MAP4 affects microtubule properties and cell cycle progression. J Cell Sci 114, 2879–2887 (2001).

5. Vasquez, R. J., Gard, D. L. & Cassimeris, L. Phosphorylation by CDK1 regulates XMAP215 function in vitro. Cell Motil Cytoskeleton 43, 310–321 (1999).

6. Ookata, K. et al. Cyclin B interaction with microtubule-associated protein 4 (MAP4) targets p34cdc2 kinase to microtubules and is a potential regulator of M-phase microtubule dynamics. J Cell Biol 128, 849–862 (1995).

7. Gergely, F., Draviam, V. M. & Raff, J. W. The ch-TOG/XMAP215 protein is essential for spindle pole organization in human somatic cells. Genes Dev 17, 336–341 (2003).

8. Hunt, T. On the regulation of protein phosphatase 2A and its role in controlling entry into and exit from mitosis. Adv Biol Regul 53, 173–178 (2013).

9. Krek, W. & Nigg, E. A. Mutations of p34cdc2 phosphorylation sites induce premature mitotic events in HeLa cells: evidence for a double block to p34cdc2 kinase activation in vertebrates. EMBO J 10, 3331–3341 (1991).

10. D’Angiolella, V., Palazzo, L., Santarpia, C., Costanzo, V. & Grieco, D. Role for non-proteolytic control of M-phase-promoting factor activity at M-phase exit. PLoS One 2, e247 (2007).

11. Oh, J. S., Susor, A. & Conti, M. Protein tyrosine kinase Wee1B is essential for metaphase II exit in mouse oocytes. Science 332, 462–465 (2011).

12. Vassilev, L. T. et al. Selective small-molecule inhibitor reveals critical mitotic functions of human CDK1. Proc Natl Acad Sci U S A 103 (2006).

13. Visconti, R. et al. The Fcp1-Wee1-Cdk1 axis affects spindle assembly checkpoint robustness and sensitivity to antimicrotubule cancer drugs. Cell Death Differ. 22, 1551–1560 (2015).

14. Serpico, A. F., D’Alterio, G., Vetrei, C., Della Monica, R., Nardella, L., Visconti, R. & Grieco, D. Wee1 rather than Plk1 is inhibited by AZD1775 at therapeutically relevant concentrations. Cancers 11, 819 (2019).

15. Serpico, A. F. & Grieco, D. Recent advances in understanding the role of Cdk1 in the Spindle Assembly Checkpoint. F1000Res 9, F1000 Faculty Rev-57 (2020).

16. Silljé, H. H. W. & Nigg, E. A. Purification of mitotic spindles from cultured human cells. Methods 38, 25–28 (2006).

17. David, A. F. et al. Augmin accumulation on long-lived microtubules drives amplification and kinetochore-directed growth. J Cell Biol 218, 2150–2168 (2019).

18. Thawani, A. et al. The transition state and regulation of γ-TuRC-mediated microtubule nucleation revealed by single molecule microscopy. Elife 9, e54253. (2020).

19. Solomon, M. J., Lee, T. & Kirschner, M. W. Role of phosphorylation in p34cdc2 activation: identification of an activating kinase. Mol Biol Cell 3, 13–27 (1992).

20. Novák, B. & Tyson, J. J. Mechanisms of signalling-memory governing progression through the eukaryotic cell cycle. Curr Opin Cell Biol. 69, 7–16 (2021).

21. Pomerening, J. R., Sontag, E. D. & Ferrell, J. E. Jr. Building a cell cycle oscillator: hysteresis and bistability in the activation of Cdc2. Nat Cell Biol. 5, 346–351 (2003).

22. Ookata, K. et al. MAP4 is the in vivo substrate for CDC2 kinase in HeLa cells: identification of an M-phase specific and a cell cycle-independent phosphorylation site in MAP4. Biochemistry 36, 15873–15883 (1997).

23. Charrasse, S., Lorca, T., Dorée, M. & Larroque, C. The Xenopus XMAP215 and its human homologue TOG proteins interact with cyclin B1 to target p34cdc2 to microtubules during mitosis. Exp Cell Res 254, 249–256 (2000).

24. Kwon, Y. G., Lee, S. Y., Choi, Y., Greengard, P. & Nairn, A. C. Cell cycle-dependent phosphorylation of mammalian protein phosphatase 1 by cdc2 kinase. Proc Natl Acad Sci U S A 94, 2168–2173 (1997).

25. Meadows, J. C. et al. Spindle checkpoint silencing requires association of PP1 to both Spc7 and kinesin-8 motors. Dev Cell 20, 739–750 (2011).

26. Visconti, R., Palazzo, L., Della Monica, R. & Grieco, D. Fcp1-dependent dephosphorylation is required for M-phase-promoting factor inactivation at mitosis exit. Nat Commun 3, 894 (2012).

27. Woodruff, J. B., et al. The centrosome is a selective condensate that nucleates microtubules by concentrating tubulin. Cell 169, 1066–1077 (2017).

28. Yim, H. & Erikson R. L. Cell division cycle 6, a mitotic substrate of polo-like kinase 1, regulates chromosomal segregation mediated by cyclin-dependent kinase 1 and separase. Proc Natl Acad Sci U S A 107, 19742–19747 (2010).

29. Schmidt, A. K. et al. The p53/p73 - p21^CIP1^ tumor suppressor axis guards against chromosomal instability by restraining CDK1 in human cancer cells. Oncogene 40, 436–451 (2021).

